# Identifying distinct neural features between the initial and corrective phases of precise reaching using AutoLFADS

**DOI:** 10.1101/2023.06.30.547252

**Authors:** Wei-Hsien Lee, Brianna M Karpowicz, Chethan Pandarinath, Adam G. Rouse

## Abstract

Many initial movements require subsequent corrective movements, but how motor cortex transitions to make corrections and how similar the encoding is to initial movements is unclear. In our study, we explored how the brain’s motor cortex signals both initial and corrective movements during a precision reaching task. We recorded a large population of neurons from two male rhesus macaques across multiple sessions to examine the neural firing rates during not only initial movements but also subsequent corrective movements. AutoLFADS, an auto-encoder-based deep-learning model, was applied to provide a clearer picture of neurons’ activity on individual corrective movements across sessions. Decoding of reach velocity generalized poorly from initial to corrective submovements. Unlike initial movements, it was challenging to predict the velocity of corrective movements using traditional linear methods in a single, global neural space. We identified several locations in the neural space where corrective submovements originated after the initial reaches, signifying firing rates different than the baseline before initial movements. To improve corrective movement decoding, we demonstrate that a state-dependent decoder incorporating the population firing rates at the initiation of correction improved performance, highlighting the diverse neural features of corrective movements. In summary, we show neural differences between initial and corrective submovements and how the neural activity encodes specific combinations of velocity and position. These findings are inconsistent with assumptions that neural correlations with kinematic features are global and independent, emphasizing that traditional methods often fall short in describing these diverse neural processes for online corrective movements.

**Significance Statement:** We analyzed submovement neural population dynamics during precision reaching. Using an auto- encoder-based deep-learning model, AutoLFADS, we examined neural activity on a single-trial basis. Our study shows distinct neural dynamics between initial and corrective submovements. We demonstrate the existence of unique neural features within each submovement class that encode complex combinations of position and reach direction. Our study also highlights the benefit of state-specific decoding strategies, which consider the neural firing rates at the onset of any given submovement, when decoding complex motor tasks such as corrective submovements.

## Introduction

Classic approaches to studying neural control of movements probe a small fraction of the behaviors critical to daily life. This limitation is often analytical, as standard approaches for interpreting neural activity rely on averaging responses across repeated behavioral trials (Pruszynski and Zylberberg, 2019). While trial-averaging helps provide robustness to highly variable neuronal responses on individual trials, it requires studying behaviors with repeated trial structure and high similarity across trials. This strategy becomes limiting for studying the neuronal processes of behaviors of increased complexity. If we wish to study sensorimotor control with the richness and complexity of natural behaviors, we must characterize neural systems on a single-trial level to understand how populations of neurons take in new sensory information combined with previous motor errors to plan, update, and generate movements.

Although common in natural reaching behavior, neuronal processes for corrective movements are understudied. During precise reaches, subjects often generate corrective movements to compensate for sensorimotor noise by incorporating visual and sensory feedback (Abrams et al., 1990; Sainburg et al., 1999; Elliott et al., 2010). In a traditional velocity-encoding model of neural activity (Moran and Schwartz, 1999), one would expect activity to return to a baseline state during the low-speed period and then encode the smaller corrective movements with similar tuning but smaller amplitude changes in firing rate with separate and additive tuning for posture (Wang et al., 2007). However, the temporal dynamics of the neural population may instead depend on the initial reaching movement and not return to a common baseline state (Albert et al., 2020). Additionally, corrective movements may not simply encode as smaller initial movements, but they may instead have distinct neural activity patterns produced by subpopulations of individual neurons (Evarts et al., 1983). These differing activity patterns observed across submovement types in the high-dimensional neural data may still be summarized with a much lower-dimensional representation but using more dimensions than initial movements. A more complete description of motor encoding requires examining not just initial, instructed movements but also how brain areas represent online corrections of movement.

When a subject reaches for but misses a target due to motor error, the corrective submovements are heterogeneous. Correction may be needed from many different postures and require movement in any direction, necessitating a unique, state-dependent correction from an initial reach. This behavioral variability may reflect greater heterogeneity in the underlying neural processes not fitting neatly into a single kinematic encoding scheme such as a single velocity- or force-based space. Analyzing the associated neural data via trial-averaging would require assumptions about which corrective behaviors lead to similar neural activity and risks obscuring the neural processes of interest. Thus, single-trial neural analysis strategies are required to uncover the neural patterns of corrective submovements. Since current experimental methods only record from a small sampling of neurons and neuronal firing rates often have stochastic features, raw, single-trial estimated firing rates can be challenging to interpret.

However, most movements including corrective movements tend to include discrete submovements (Paninski et al., 2004; Fishbach et al., 2007; Hatsopoulos et al., 2007). The submovements are further defined by condition-invariant neural activity (Kaufman et al., 2016; Kobak et al., 2016) and the distribution of peak speeds . Our recent work identified consistent and distinct peaks in speed that define separate initial and corrective submovements with a low-speed period almost always occurring between each submovement for visually guided precision reaching (Rouse et al., 2022). These submovements correspond to cyclic and asynchronous rise and fall of firing rates across the neural population that provide information about when each peak in speed occurs. Thus, this structure of submovements raises the possibility that the latent state of the neural population on any single trial and within each submovement may influence how each corrective movement is signaled. Here, we analyze data from a precision reaching task to understand how neural firing rates encode various corrective submovements and how reach velocity predictions can be improved.

An additional technical challenge for studying corrective movements is neural recording interface instability, such as movements of the electrode tip relative to the recorded neurons, which prevents consistent recording from the exact same neurons across recording sessions (Chestek et al., 2011; Fraser and Schwartz, 2012; Perge et al., 2013). These instabilities limit the total number of trials with a given behavior that we can observe for any given neuron. Thus, for behaviors with a high degree of heterogeneity, such as corrective movements, it becomes challenging to gather sufficient data within a single recording session to characterize the neural and behavioral spaces of interest. As a result, any analysis of highly variable corrective movements will also require analysis techniques that “stitch” recording sessions together into a common representation of neural activity (Turaga et al., 2013; Nonnenmacher et al., 2017; Pandarinath et al., 2018b; Degenhart et al., 2020).

Here we address the limitations in characterizing the neuronal processes underlying corrective movements using deep-learning–based single-trial analytic methods. This allows a novel comparison of neuronal dynamics during not only initial but also corrective movements. We examined neuronal population activity recorded from primary motor cortex during a precision-movement task that elicited a range of initial and corrective movements. We apply a previous approach, AutoLFADS (Keshtkaran et al., 2022), to uncover stable representations of neuronal population dynamics and now aggregation of data from many sessions. Importantly, we are able to identify the most neurally similar movements across many sessions and then examine what is shared in their kinematics to better infer how the brain organizes its repertoire of movements.

## Materials and Methods

### Experimental preparation

All procedures for the care and use of nonhuman primates followed the *Guide for the Care and Use of Laboratory Animals* and were approved by the University Committee on Animal Resources at the University of Rochester, Rochester, New York. Two male rhesus monkeys, P and Q (weights 11 and 10 kg, ages 7 and 6 years old, respectively), performed the task in the study.

### Behavioral analysis

The monkeys performed a center-out task using an 18 cm–long cylindrical rod as a joystick to control a cursor on a 24" LCD display. The joystick could move freely with little resistance. The cursor on the screen represented the joystick position which freely moved within a 1000-by-1000-pixel workspace. A small cross symbol served as the cursor to indicate a single point in the workspace. Customized software sampled the joystick data, refreshed the scene, and saved the cursor position and trial event times at 100Hz.

We recorded 12 sessions of neural recordings for monkey P and 12 sessions for monkey Q over a time span of 2 months and analyzed the recorded neural signals using AutoLFADS. One session for monkey Q was excluded due to greater variation in the behavior likely due to reduced motivation by the animal that AutoLFADS could not resolve. The two monkeys successfully completed a total of 10,958 (monkey P) and 8737 (monkey Q) trials across recording sessions. The task as well as analysis of behavior and condition-invariant neural activity have been described previously (Rouse et al., 2022). Briefly, in the task trials, after obtaining and holding the cursor for 300-500 ms in a center target with a radius of 75 pixels, the monkey would reach from the center to 1 of 8 instructed directions, equally spaced by 45° around the origin as shown in Fig. 1A. Each of those peripheral directions included one of three different sizes defined as large, shallow, and narrow targets. Different target sizes were introduced into the task to vary difficulty. The shallow and narrow targets induced more frequent corrective movements since they were smaller. Our present analysis, however, includes all initial and corrective movements since we did not identify significant differences in the corrective movements dependent on the target size, aside from their frequency. The monkey would receive a liquid reward once he successfully made a complete hold of 500-600 ms with the joystick within the peripheral target. The required initial and final hold times for each trial were randomly and uniformly sampled from within the given range.

**Figure 1.**
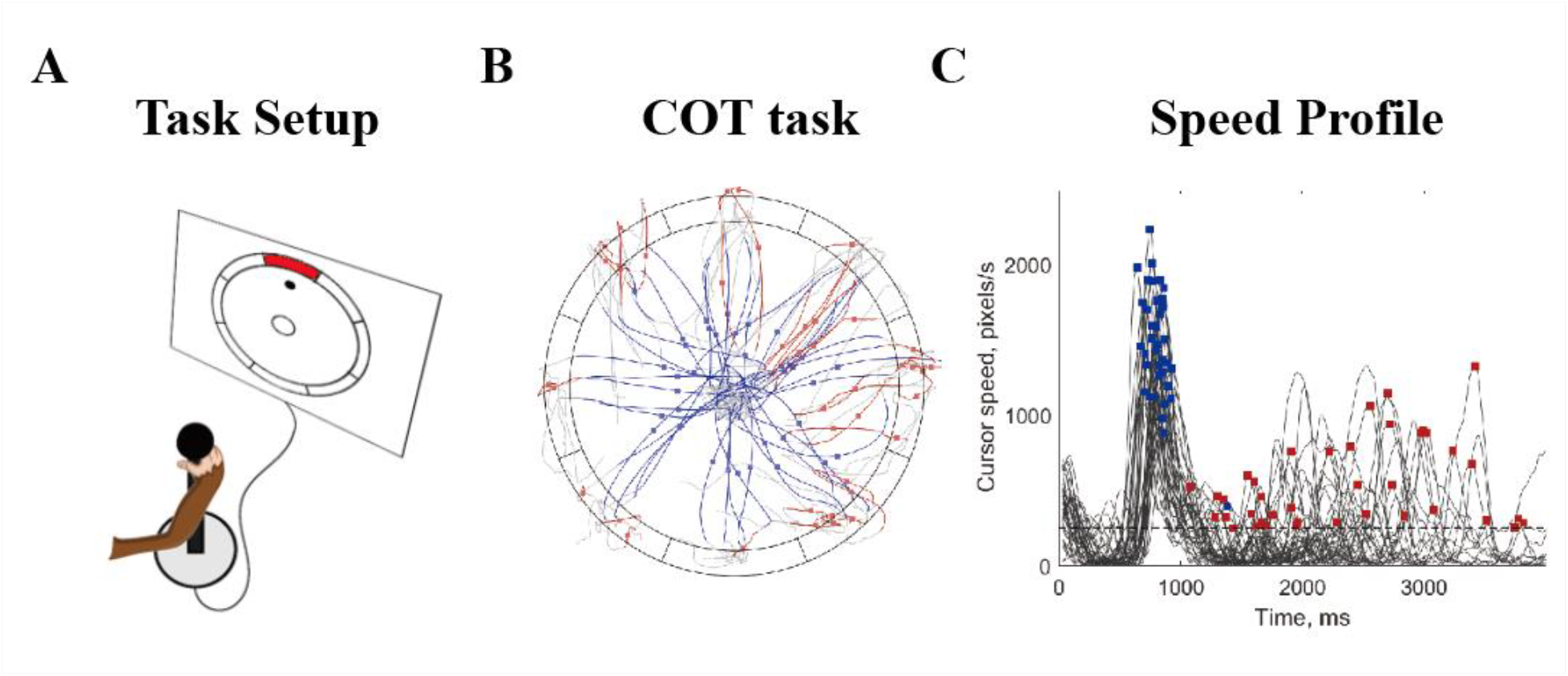
Behavioral task schematic. A) Behavioral task scheme showing the control and display of cursor. B) The schematic for the center-out task displays the targets arranged in eight uniformly distributed outer blocks, with the initial reaching movements illustrated in blue and the corrective reaching movements in red. C) Cursor speed profile of the movement trials over time, where initial (blue) and corrective (red) submovement speed peaks are identified with squares. A distribution of the peak speeds is provided in Figure 1-1.

The behavioral task schematic and cursor speed profile for a subset of trials are shown in Figure 1. Fig. 1B demonstrates the reach trajectories for the shallow targets. The monkey would acquire the center target and then reach towards 8 peripheral targets (the shallow outer boxes). Movement trajectories of the initial reach submovements are indicated in blue, and the speed peaks of each trial are marked as blue squares. Corrective submovement trajectories are shown in red, and the speed peaks of each trial are marked with red squares. Gray lines connect the rest of a trial before, between, or after submovements with a speed peak. Fig. 1C shows the cursor speed profile over time throughout the trials. The initial peak speeds (blue boxes) and corrective peak speeds (red boxes) are marked as the annotation points across the speed profile plot. We calculated cursor speed offline by filtering cursor position with a 10 Hz low-pass first-order Butterworth filter (bidirectionally for zero phase lag). We used a peak-finding algorithm to identify speed peaks (findpeaks function in Matlab (Mathworks, 2021)) to define both initial and corrective submovements. Cursor speed peaks were identified when the following criteria were met: i) the peak speed was greater than 250 pixels/s and ii) the peak had a prominence (the height difference between the peak and the larger of the two adjacent troughs) of at least 50% of the absolute height of the peak. Submovements within a trial are defined as the surrounding time window of 400 ms before and 200 ms after cursor speed peaks. We defined the initial speed peaks to be the first submovements which ended at least 150 pixels from the center, and any small movements before were excluded from further analysis. Speed peaks following the initial speed peak were defined as corrective submovements, and any corrective submovements occurring only within the peripheral target were excluded to focus analysis on submovements made to successfully reach towards the target. These corrective speed peaks were significantly smaller in magnitude with only 5% and 13% of submovements overlapping with the slowest initial submovements in monkey P and Q, respectively (Ext. Fig. 1-1). Over all characterized submovements, there were 6464 and 3728 corrective submovements identified for monkeys P and Q, respectively. Across all trials, 17.5% (P) and 20.2% (Q) of trials required one additional corrective submovement, and 14.2% (P) and 8.7% (Q) of trials required more than one correction after an initial movement.

**Figure 1-1.**
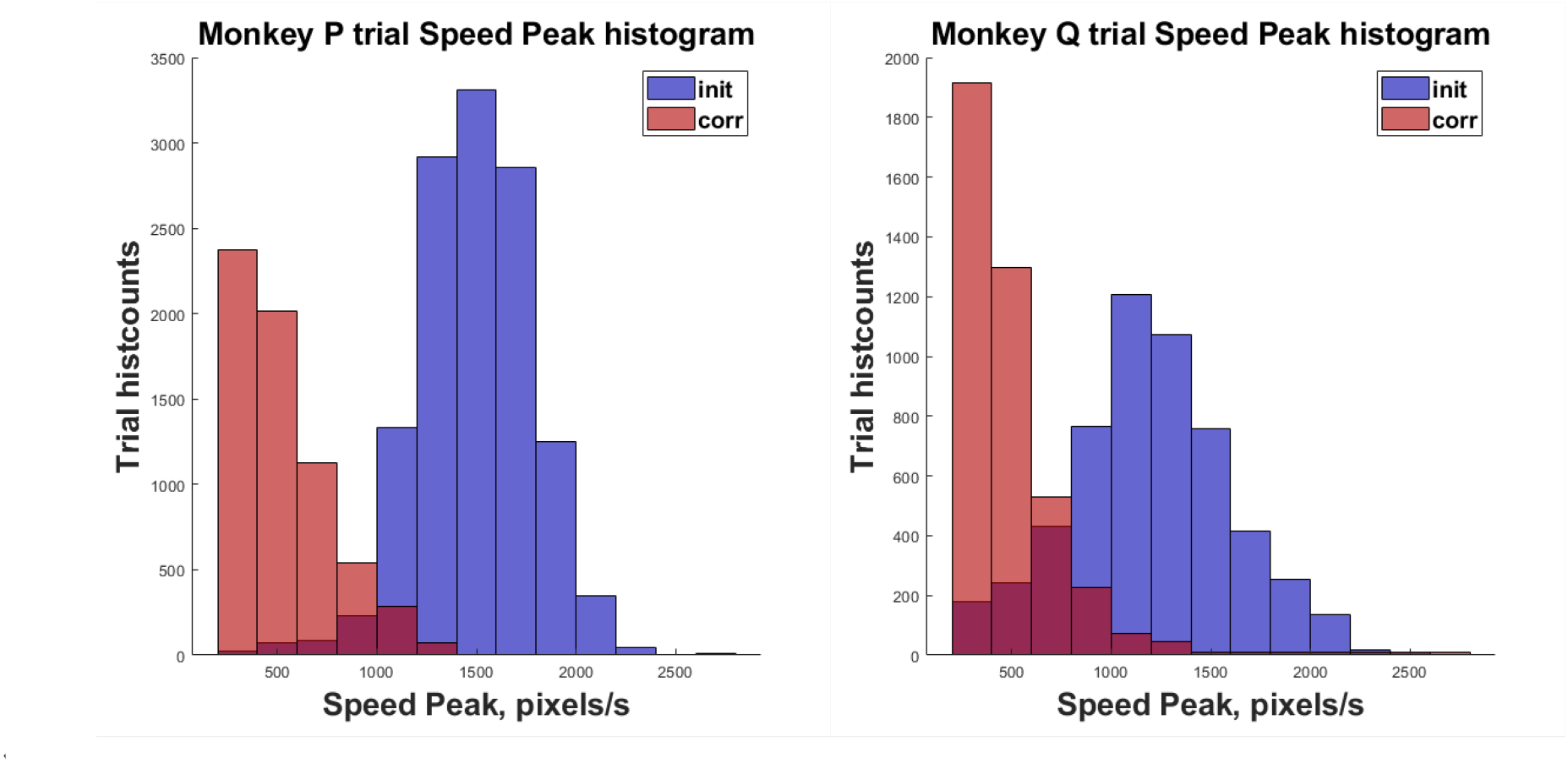
Histogram of trials speed peak of the two monkeys. The distribution in speed peaks for the two monkeys over all stitched sessions for both initial (blue) and corrective (red) submovements. Both animals had little overlap between the magnitude of initial and corrective submovements but Monkey Q had more trials (13% of the total trials) with overlap speed peak between initial and corrective movement then Monkey P (5% of the total trials).

### Neural Recordings

Floating microelectrode arrays (MicroProbes for Life Science) were implanted in the anterior bank of the central sulcus in each monkey to record signals from primary motor cortex (M1) as described previously (Mollazadeh et al., 2011; Rouse and Schieber, 2016; Rouse et al., 2022). Recordings were collected from six 16-channel arrays implanted in M1 for monkey P and two 32-channel arrays plus one 16-channel array implanted in M1 for monkey Q. Plexon MAP and Blackrock Cerebus data acquisition systems were used to manually threshold the spiking activities and collect spike-snippets and spike times which were then offline sorted using a custom, semi-automated process (Rouse et al., 2022).

### AutoLFADS and ‘Stitching’ process

We analyzed all neural and cursor data by time aligning to the identified speed peaks used to define each submovement. We preprocessed the raw neural data by calculating the trial perimovement time histogram (PMTH) in 20 ms sampling bins. We then analyzed the neural data for each submovement from 400 ms before peak speed to 200 ms after peak speed.

We then applied the deep-learning based technique AutoLFADS to identify latent factor representations of neuronal population dynamics (Keshtkaran et al., 2022). In brief, AutoLFADS is a method for automated hyperparameter tuning of latent factor analysis via dynamical systems (LFADS) models.

LFADS is a deep learning approach that uses recurrent neural networks to model the spatiotemporal patterns of neural data (Pandarinath et al., 2018b). Specific values pertaining to our AutoLFADS training can be found in Ext. Table 2-1.

**Table 2-1.**
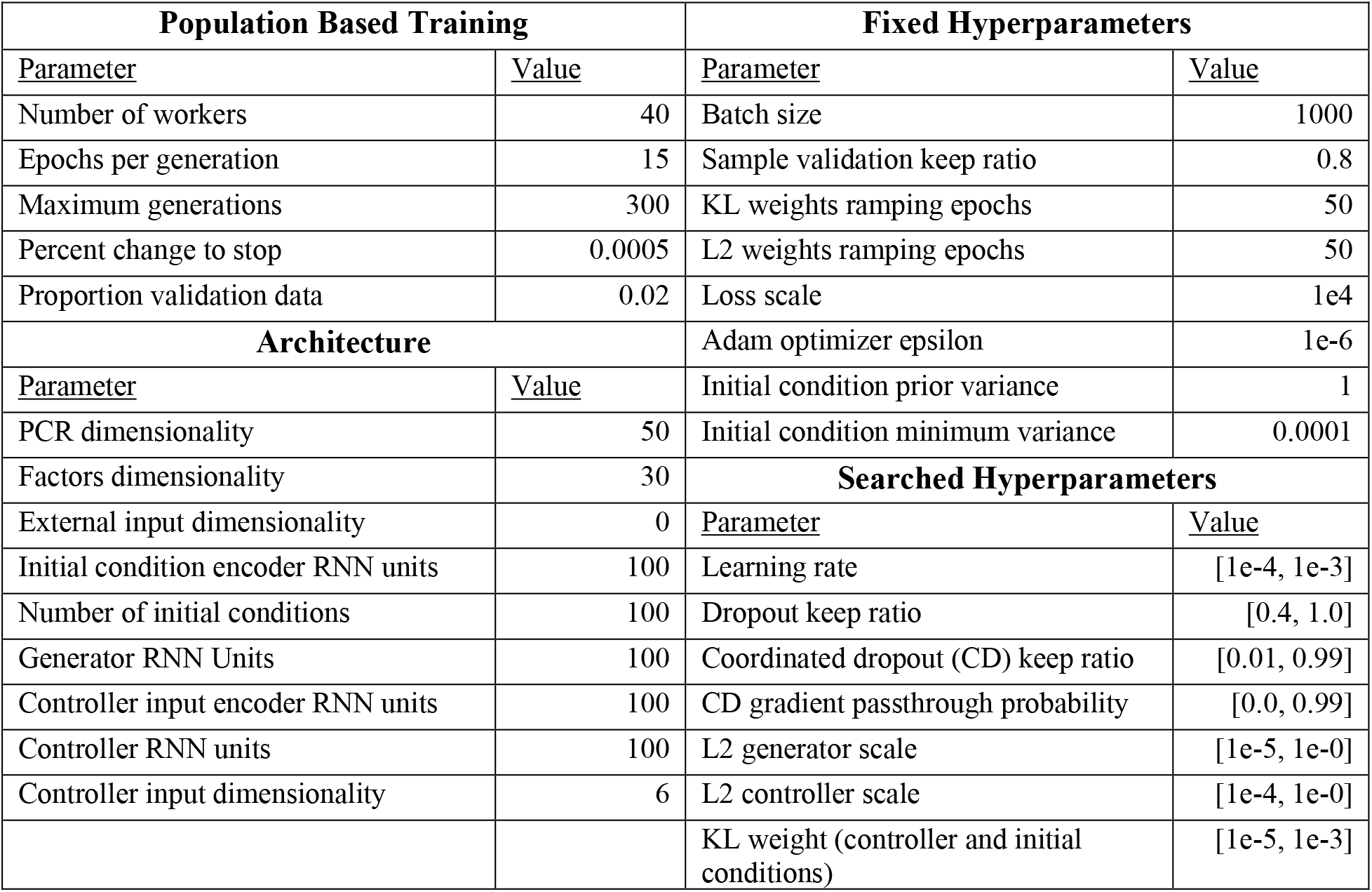
AutoLFADS Architecture, Hyperparameters, and Search Settings.

We combined the automated hyperparameter optimization of AutoLFADS with LFADS model *stitching*, which has been previously shown to identify latent factors that are consistent with behavior across months, allowing data aggregation from multiple recording sessions (Pandarinath et al., 2018b). To execute stitching, multiple sessions of preprocessed neural data were sent to a configured AutoLFADS model using the pre-session ‘read-in’ matrices. Each read-in matrix is fixed with weights W^input^, which are learned from a principal component regression (PCR) that maps the trial-averaged firing rates from each individual session to the shared principal components across all sessions. Passing the spiking data through these read-in matrices provides a set of input factors. Then the input factors were fed into a single encoder, generator and factor matrix W^fac^ shared across sessions to get output factors within a common latent state-space across all sessions. Using this across-sessions shared latent space, we then obtained factors corresponding to each session for use in downstream analysis. This process is summarized in Ext. Fig. 2-1.

**Figure 2-1.**
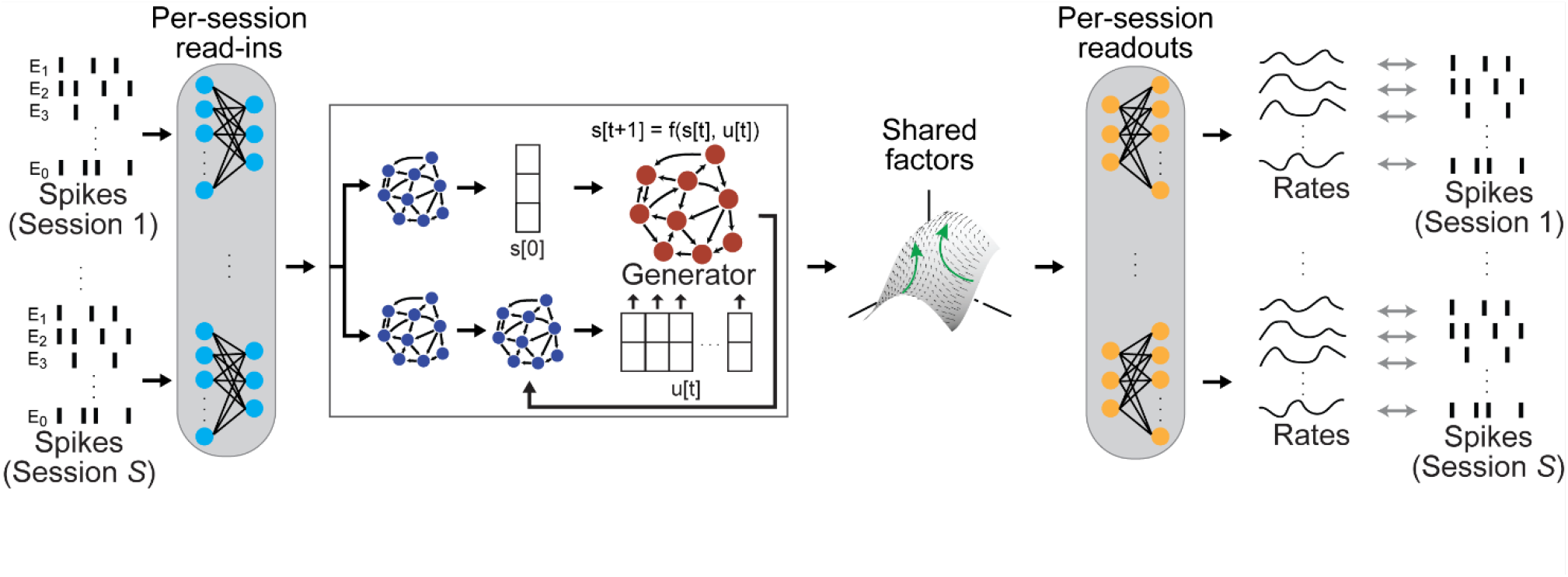
Multisession LFADS schematic. LFADS stitching from Pandarinath et al. (2018) allows a single latent space to be used to describe many sessions of neural data simultaneously. The method initializes a read-in matrix for each session using principal component regression (PCR) to obtain weights that map the condition-averaged data from each session to the shared principal components of all sessions. These read-in matrices are held fixed during training, encouraging the data being input to the LFADS model to approximate a shared space. A single LFADS model is trained to learn the shared latent space across all sessions. These shared factors are then mapped back out to the neural data from each session using a per-session read-out matrix that is initialized using the PCR weights but allowed to train with the rest of the model.

### Quantitative analysis on measuring correlation between LFADS factors and kinematics

We used a total of four different analysis methods to verify the validity of our LFADS factors representation, examine the neural space, and decoded neural activity. For visualization, both a linear— principal component analysis (PCA) — and a non-linear—t-distributed stochastic neighbor embedding (t-SNE)— method were used. Two other methods—ridge regression and a novel state dependent decoder—were used to test how well movement could be predicted from decoded neural activity.

### Principal Component Analysis

PCA (pca function in Matlab (Mathworks, 2021)) was used to reduce the dimensionality of the LFADS factors for visualization while maintaining the most variance in the neural data. PCA imposes a minimal number of assumptions about the underlying structure of data and reveals the dimensions that explain most of the variance. We projected the LFADS factors into a single PCA subspace across all sessions as a function of time to create neural trajectories and reveal the neural dynamics for given submovements.

The neural states at the start of each submovement were also tested for what fraction of variability was accounted for by submovement types (initial vs. corrective) and starting cursor position. We calculated the pairwise distances between initial and corrective submovement trials within the LFADS factor space on the first time point to quantitatively describe the discriminative ability between initial and corrective submovements. Similarly, we calculated the pairwise neural distance between the 4 opposite positions in the physical work space—up vs. down, left vs. right, up/right vs. down/left, and up/left vs. down/right— to measure the discriminability between positions expected to have the largest difference in neural activity.

### t-Distributed Stochastic Neighbor Embedding

To better identify neurally similar movements within the high-dimensional neural data and visualize the data in a low-dimensional subspace, we used t-SNE, a non-linear dimensionality reduction tool. We mapped 30 dimensions of LFADS factors from individual trials into a 2D space using the built-in t-SNE toolbox in Matlab. Data fed into t-SNE consisted of LFADS output with time value from 100ms before and 100ms after peak speed sampled every 20ms. We resized the stitched latent factors by stacking the time and factors dimensions, from [trial × time × factor] to [trial ×(time × factor)] to identify trials with similar factors across time. We performed t-SNE with the Barnes-Hut algorithm using the Euclidean distance metric with perplexity set to 50 and learning rate to 500 (Wattenberg et al., 2016). The t-SNE output was set to two dimensions to introduce a simple and clear view of data structure. -.

The t-SNE map, which identifies submovements with similar neural activity, was then subdivided to analyze corresponding movement trajectories in the two-dimensional physical workspace. For identifying and visualizing similar individual trials into clusters, we utilized *k*-means nearest neighbors clustering on the t-SNE space of LFADS factors for initial and corrective submovements separately. The *k*-means cluster number was determined by calculating the sum square of distance to the centroids for increasing number of clusters. By visual inspection, the elbow method was used to identify when additional clusters would not significantly improve the average movements’ distance from the centroid. The kinematics, approximated by their straight-line vector, could then be examined for submovements within each cluster.

### Ridge Regression

We used ridge regression to quantify the correlation between neural activity and the submovement kinematics. We refer to this process of mapping between neural data and behavior as *decoding.* To allow comparison between neural representations, we used ridge regression to map between either the smoothed firing rates or the AutoLFADS latent factors and the submovement kinematics.

Ridge regression was implemented as in Scikit-Learn (Pedregosa et al., 2011), where the regression model solves for some set of weights W that minimize the linear least squares loss in the following equation:

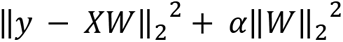

where X is the neural data, *y* is the behavioral cursor velocity data to predict, ||.|| ^2^ is the squared L2 norm, and ɑ is the L2 penalty applied to the weights.

Prior to training the ridge regression model, we applied a lag to the data such that the neural activity preceded movement by 100ms (Ext. Fig. 2-2 A,C). The ridge regression model was trained using five fold cross-validation with an L2 regularization value of ɑ = 1.0 (Ext. Fig. 2-2 B,D). Final decoding accuracy was reported as the average velocity R^2^ of the held-out folds for within-day decoders, or as the velocity R^2^ for held-out data (sessions or trials).

**Figure 2-2.**
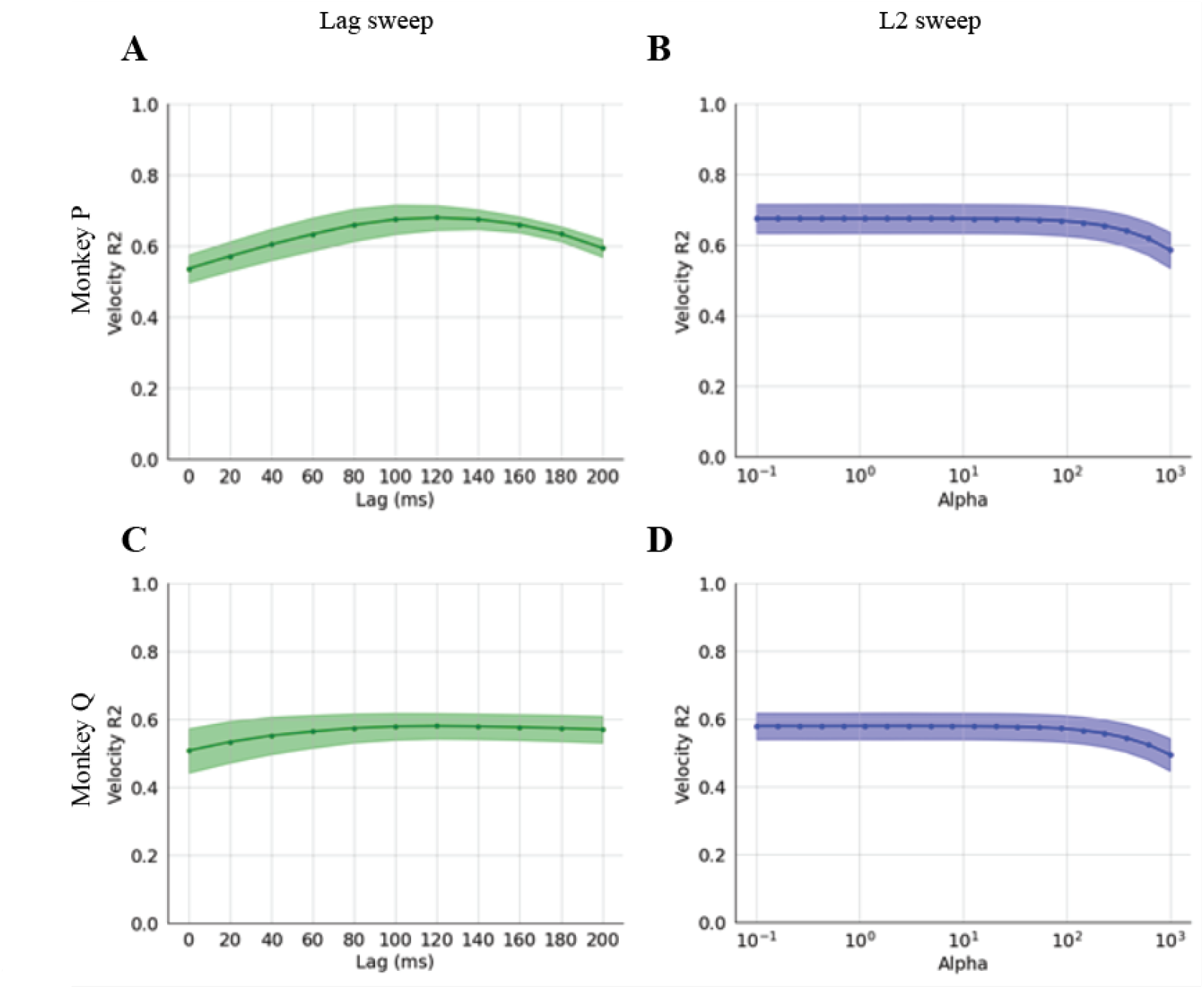
Grid search of decoding lag and alpha value to determine ridge-regression parameters. The predicted velocity R^2^ from smoothed spikes for Monkey P sessions at (a) different lags and (b) different regularization alpha values. Solid line shows the average across all datasets and the shaded region is the standard deviation. (c-d) same as a-b for Monkey Q. Based on this grid search, a lag of 100 ms and alpha of 1 was used for all further analysis.

### State-dependent decoder

Due to apparent differences in neural state at the start of submovements for corrective submovements (see Results below), a discrete state-dependent linear decoder was used to measure improvements in decoding performance when classified by different states based on the starting neural state. Different states were defined by *k*-means clustering of the t-SNE space of LFADS factors at a timepoint before the corrective submovements (340 ms before peak speed). The number of clustered states was varied from 1 to 14 in the analysis. After performing *k*-means clustering on the test data, a separate ridge regression was then trained on each *k*-means cluster. For decoding, each test trial was assigned to its nearest cluster centroid by its neural activity 340 ms before peak speed. The ridge regression coefficients for the given assigned cluster were then used to transform the neural activity into predicted movement.

Regressions were performed using 5-fold cross-validation. All trials were divided into 5 equal testing groups with the remaining 80% of data used for training the prediction for each test group. Global sum of squares residuals and total sum of squares total were manually calculated and summed across the states to return the variance predicted in the kinematic dataset and the total variance. The coefficient of determination (R^2^) for the state-dependent decoders was then calculated with the variance values.

Bootstrap testing by selecting trials with replacement for 1000 iterations was used to estimate 95% confidence intervals.

## Results

### Stitched AutoLFADS infers consistent neural dynamics across sessions

We used AutoLFADS with stitching to obtain consistent neural representations of submovements across sessions to address the inherent constraint of limited trials available per recording session. We first evaluated AutoLFADS for its ability to consistently extract these single submovement neural population dynamics across sessions. In Fig. 2A, we visualized the stitched AutoLFADS latent factors corresponding to initial submovements from 6 representative sessions by projecting them to a 3D PCA subspace and color coding by the 8 reach directions. All subplots showed a similar neural pattern in the PCA subspace, indicating that AutoLFADS successfully identified shared latent structure across all sessions. We then used t-SNE on the neural latent factors to also show the eight initial reach directions could be consistently identified and stitched by their latent factors across sessions. In Fig. 2B, t-SNE of all sessions’ initial submovements are color coded by reach velocity vectors between 100 ms before and 100 ms after peak speed in the left subplot. In the middle and right subplot, two representative sessions were selected for visualization (colored by reach velocity vectors); all other sessions of submovements were colored as gray. Together, Figs. 2A and 2B show AutoLFADS generates consistent latent factors for the initial reaching submovements and stitching creates a common neural space across days corresponding to the 8 reach directions.

**Figure 2.**
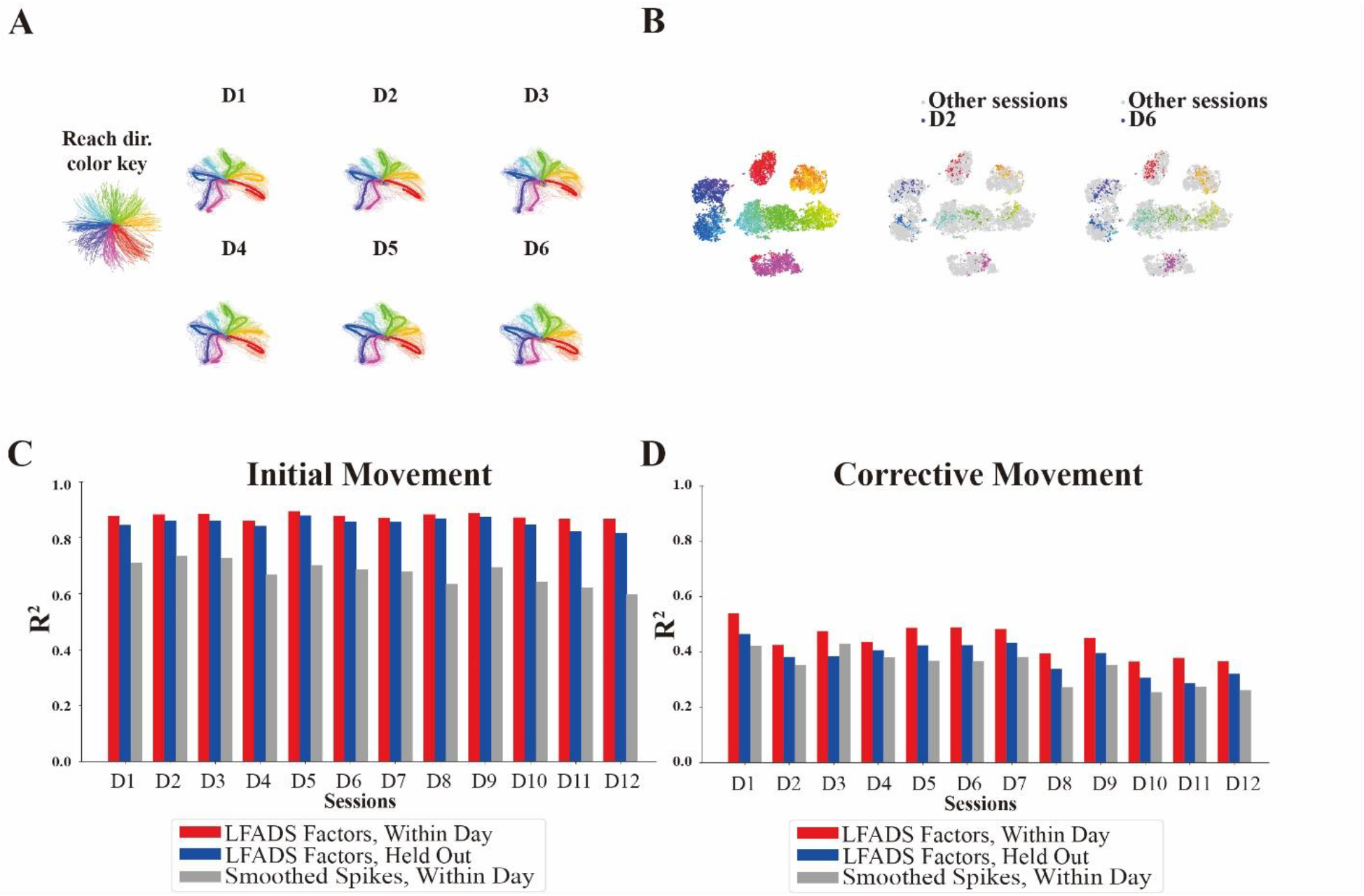
AutoLFADS and stitching enables analysis of cross-session neural data. A) Neural trajectories for each single session are shown in the subplots in the first three PC space of LFADS factors. The single trials in each session are presented as thin lines in the subplot and average trajectories for 8 different directions are presented in bold colored lines. The color key for each reaching direction is shown on the left. B) t-SNE plot of initial submovements across all sessions (left) and with 2 representative single sessions highlighted (right), color coded by movement direction. C,D) Correlations between kinematic and latent factors of both held-in/held-out dates for LFADS factors and held-in dates for smoothed spikes, shown separately for initial submovements (C) and corrective submovements (D) . These results indicate that inferred neural dynamics after the ‘stitching’ process can generalize to single sessions not in the original training set and also better correlate to the kinematic data than smoothed firing rates. A schematic of the AutoLFADS stitching procedure is provided in Figure 2-1. All analysis performed with time lag of 100ms and alpha of 1. The decoding R2 over various time lag and alpha values over a range of values is given in Figure 2-2. All other parameters used for AutoLFADS are provided in Table 2-1.

We further assessed how well the latent factors can capture single-trial movement variability by utilizing ridge regression. We fit the regression model from latent factors to movement velocity of initial and corrective submovements in Figs. 2C and 2D, respectively. For each set of submovements, we fit a single ridge regression model on all but one session and evaluated how well that model could decode velocity on the held-out session. For reference, we also fit a separate ridge-regression model on the smoothed spiking data from the same held-out session. The red bar for each session shows the variance explained (R²) of the behavioral predictions from LFADS factors when a regression model is trained on only that session. The blue bar for each session shows the R² of behavioral predictions from the held-out day when a regression model is trained on the LFADS factors of all other datasets. The gray bar for each session shows the R² of behavioral predictions when a regression model is trained using smoothed firing rates from the same day. For both monkeys analyzed, there was a small drop in performance from held-in to held-out R² as expected. The average drop in R² was 0.02 (monkey P) and 0.13 (monkey Q) for initial submovements and 0.06 (P) and 0.15 (Q) for corrective submovements. Even with the small decrease in R² for the held-out LFADS factors, the held-out regression was favorable compared to regression with smoothed firing rates performed on a given session. For initial submovements, both monkey P and monkey Q had significantly better R² (p<1e-8 and p=0.02, respectively, paired t-test) for held-out LFADS factors compared to smoothed spikes. For corrective submovements, LFADS factors were significantly better for monkey P (p<1e-2, paired t-test), and there was no statistical difference for monkey Q (p=0.675, paired t-test). Overall, these results indicate a significant improvement in initial decoding performance and a slight improvement or similar performance for corrective movements using AutoLFADS in both monkeys compared to smoothed firing rates. We also note that the predictions for corrective submovements yielded worse decoding performance than those for initial submovements. This indicates that there is likely more diversity in the neural encoding of corrective submovements and a single linear velocity tuning model does not describe the data as well as it does for the initial submovements.

### Corrective submovements are encoded using a combination of reach direction and location

Given that corrective submovement latent factors cannot predict movement direction as well as initial, we next examined how the neural activity for the corrective submovements related to the position and velocity of individual submovements. We defined the position of each submovement as the starting position at 100 ms before peak speed. A velocity vector of the submovement from 100 ms before to 100 ms after peak speed defined the direction and magnitude of each reach. We performed t-SNE to project the high-dimensional neural data into a 2-dimensional subspace as shown in Fig. 3, with each scatter point representing a submovement. We applied t-SNE on corrective submovements of a single session of smoothed firing rates and projected to the t-SNE space, color coded based on the position (Fig. 3A) and velocity vectors (Fig. 3B). For the smoothed firing rates t-SNE subplots, only a weak and noisy relationship with both the position and velocity vectors was observed.

**Figure 3.**
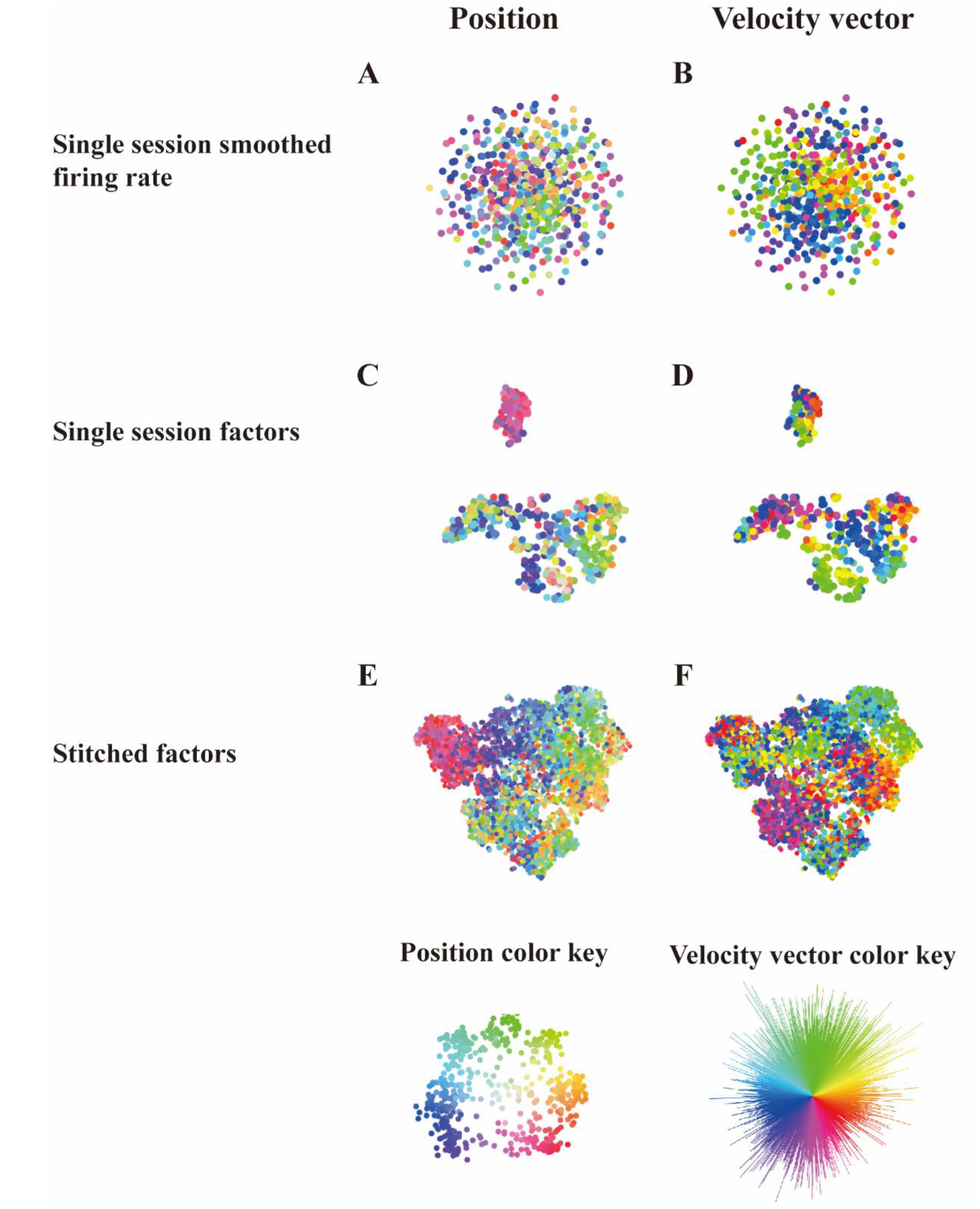
t-SNE cluster plots of corrective submovements in A/B) smoothed firing rates space C/D) latent factors space for a single session and E/F) latent factors space across all sessions. A/B) Single session t- SNE plots of corrective submovement color coded by position and velocity vector projected in smoothed firing state space. C/D) Single session t-SNE plots of corrective submovements color coded by position and velocity vector projected in latent factor space. E/F) ‘Stitched’ session t-SNE plots of corrective submovements color coded by position and velocity vector projected in latent factor space. Bottom row shows the color key for position and velocity vector color coding. Left panels are color coded for position at the onset of each submovement 100 ms before peak speed using the direction from the center for color hue and the distance from the center for saturation. Right panels are colored by reach direction using the straight-line reach direction from 100 ms before until 100 ms after peak speed. These results show that AutoLFADS factors can better identify and separate the neural patterns related to the corrective kinematics. A quantification of this improved alignment of kinematics with AutoLFADS factors compared to smoothed firing rates is provided in Figure 3-1. Further, stitching across sessions is necessary to have a large enough sample of the diverse movements to better identify structure in corrective movements.

Next, we projected LFADS factors of a single session from the stitched model into t-SNE space, again color coded on position (Fig. 3C) and velocity vector (Fig. 3D). Some distinct groupings based on kinematics are revealed but the overall relationship remains noisy and unclear. Finally, when all sessions of LFADS factors were stitched together and visualized in the t-SNE space, clearer groups and gradients based on a combination of position (Fig. 3E) and velocity vector (Fig. 3F) are observed. We performed KNN analysis on t-SNE of LFADS factors and smoothed firing rates for corrective submovements (Ext. Fig. 3-1) to calculate how similar the kinematics were for the closest neural neighbors of a given submovement in the various neural representations. The average KNN distances of both the single-session LFADS model’s factors and the stitched LFADS factors corresponding to the same single session are significantly smaller than those for single-session smoothed firing rates (p<0.05, paired t-test, Ext. Fig. 3-1), indicating LFADS factors better organized similar trials in a general neural space. Additionally, there was no significant difference between mean distance when using LFADS performed on a single session and using the stitched LFADS factors for any given session (p>0.05, paired t-test, Ext. Fig. 3-1), further indicating that LFADS factors are consistent across sessions. Interestingly, the different clusters of corrective submovements are not dominated by velocity vector or position alone but are a combined interaction between the two. For some clusters, all submovements within a dense area appear to be related to reach direction while other groups of submovements appear to be more related to a distinct location in the workspace. This provides evidence for a combined neural representation of the corrective submovements that depends on both where the submovement originates as well as the direction of the movement.

**Figure 3-1.**
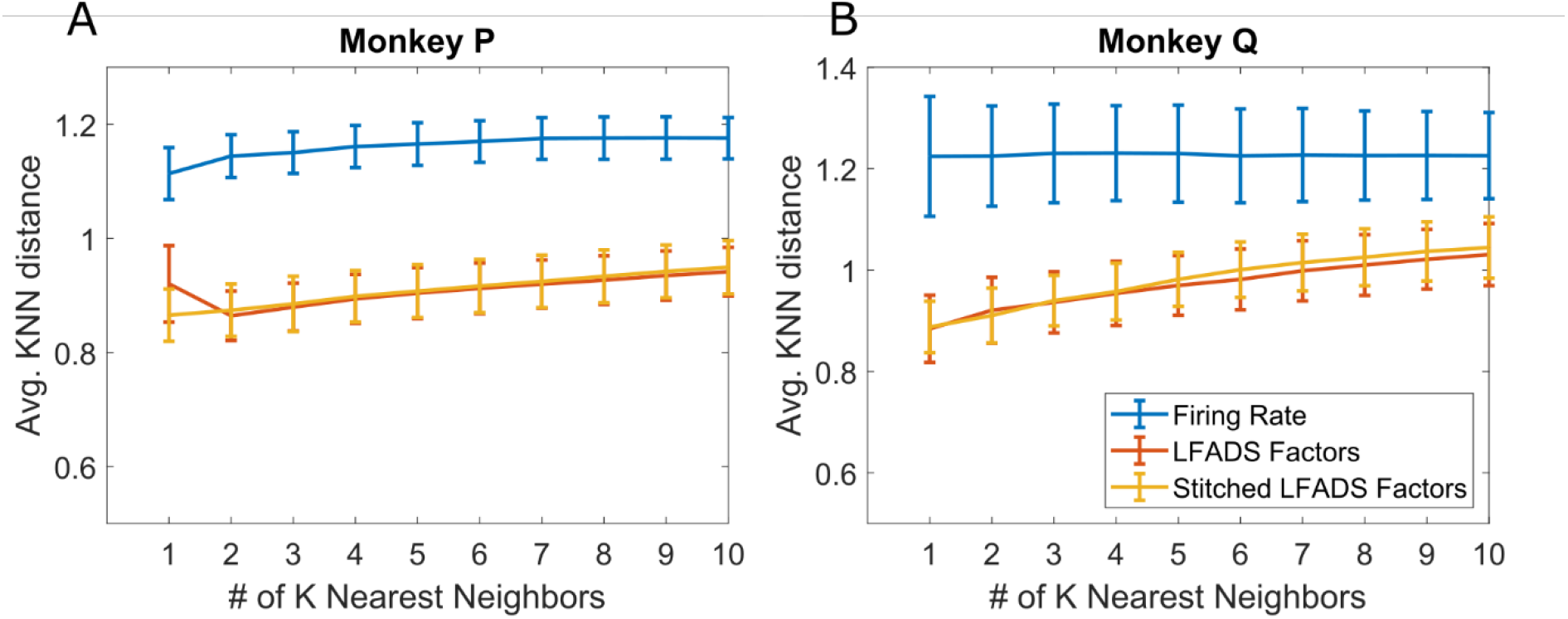
Quantification of accuracy in movement representation in the various neural spaces. The K nearest neighbors for corrective submovement firing rates from a single session, LFADS factors from a single session, and LFADS factors stitched across sessions were calculated for each submovement. The mean distance between the velocity vector for the given submovement and its K nearest neural neighbors was calculated to measure how similar the submovments were for neural data identified to be similar. Error bars are standard deviation across recording sessions. Both individual LFADS factors from a single session and stitched LFADS factors identified neural neighbors with significantly less differences in kinematics than the single-session smoothed firing rates (p<0.05, paired t-test). There was no significant difference between LFADS factors calculated for single sessions versus the stitched factors (p>0.05, paired t-test). This was true over all 1-10 K-nearest-neighbors for both monkey P (A) and monkey Q (B), demonstrating that LFADS factors more effectively organize the neural data with similar movements being closer to each other in a more cohesive neural space than smoothed firing rates.

### Distinct neural patterns associated with kinematics revealed within neural state-space

We then explored the distinct neural patterns between various submovements by examining the neural trajectories and movement trajectories for groupings identified using the t-SNE representation of neural data. In Fig. 4A, the same t-SNE plot of initial submovement LFADS factors as shown in Fig. 2B was classified by *k*-means into 8 groups. We determined 8 clusters for initial submovements, by the elbow method (Ext. Fig. 4-1), which also is expected for an 8-target task. In this case, the *k*-means classification was based only on the Euclidean distance in t-SNE neural space, not kinematic information. Fig. 4B shows the neural trajectories of initial submovements in the plane of the first two principal components. The average initial neural trajectories originate from a similar point and show 8 distinct curved trajectories in the PC plane. We then plotted the 8 groups of submovement trials separately in the kinematic space to show the corresponding movements of this neural data (Fig. 4C). The movement trajectories show good alignment with the 8 directions of the instructed targets, highlighting the strong discriminability of initial reach direction from the neural data processed with AutoLFADS. Notably, for these initial submovements in the PC plane, 6 of the 8 trajectories move towards a similar end point while the trajectories for 2 of the groups (purple and dark yellow) move to a different location in the neural space. The two trajectories with different ending locations are associated with targets in the bottom-right and bottom part of the workspace, indicating a potential posture difference for the monkey to hold the manipulandum in those two workspace locations.

**Figure 4.**
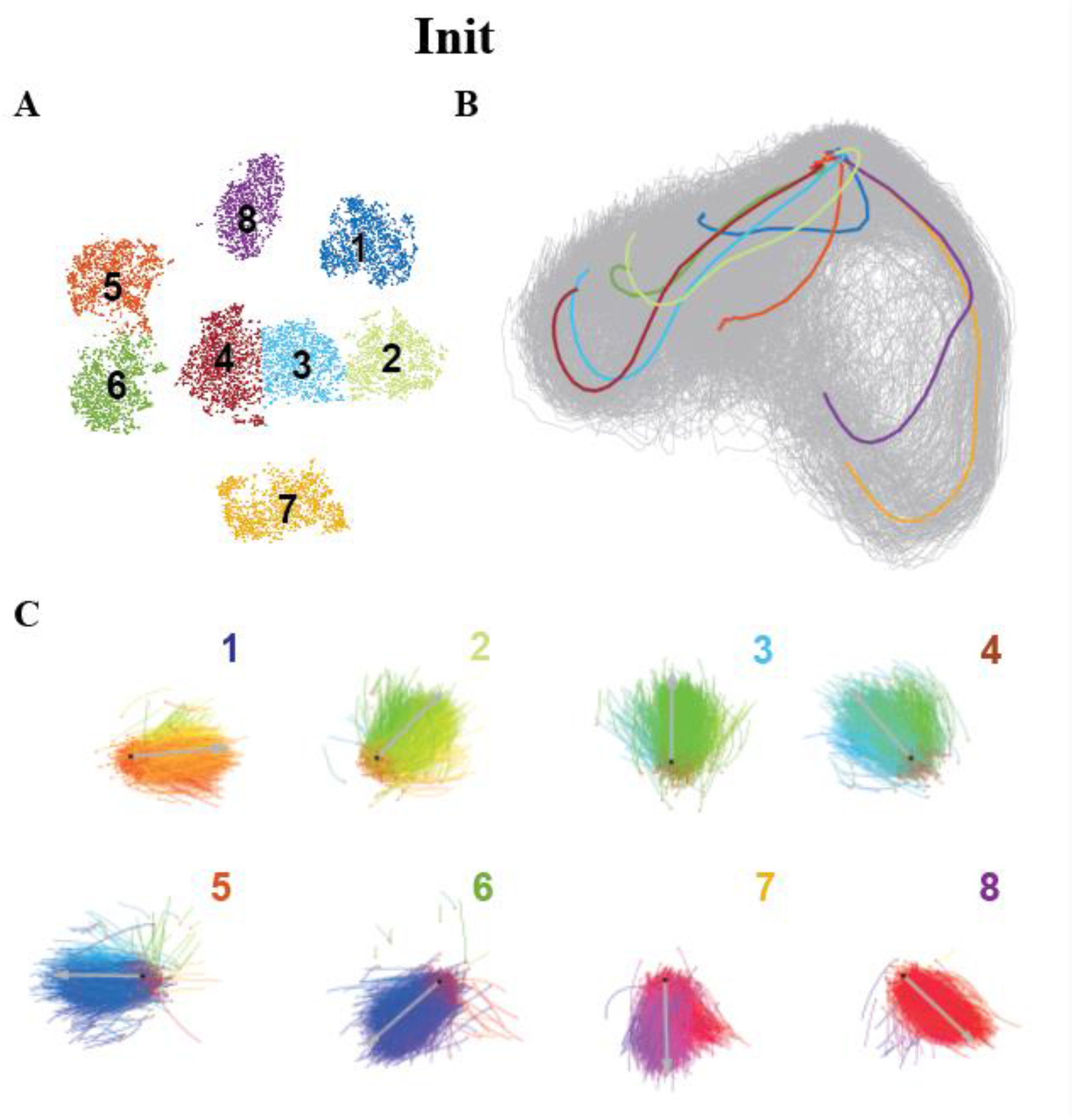
The identified organization of submovements in the neural space and their corresponding kinematics for corrective submovements. A) t-SNE plots of initial and corrective submovements AutoLFADS factors, color coded by subgroup identified with k-means clustering. We selected the number of *k*-means groups to be 8 for initial submovements using the elbow method shown in Figure 4-1. B) Neural trajectories of initial submovements factors. Average neural trajectories are color coded based on the t-SNE subgroups. All single-trial neural trajectories are colored gray in the trajectory plot. C) Movement trajectories for each subgroup identified based on the neural activity. The colored number for each subgroup corresponds to the subgroups shown in panels A&B. The movement trajectories are color coded based on velocity vector (same color key as Fig. 3). Within each subplot, the center target position was marked with a black dot, start position of each submovement was marked with red dot, and average velocity vector and relative amplitude are annotated with a gray arrow.

**Figure 4-1.**
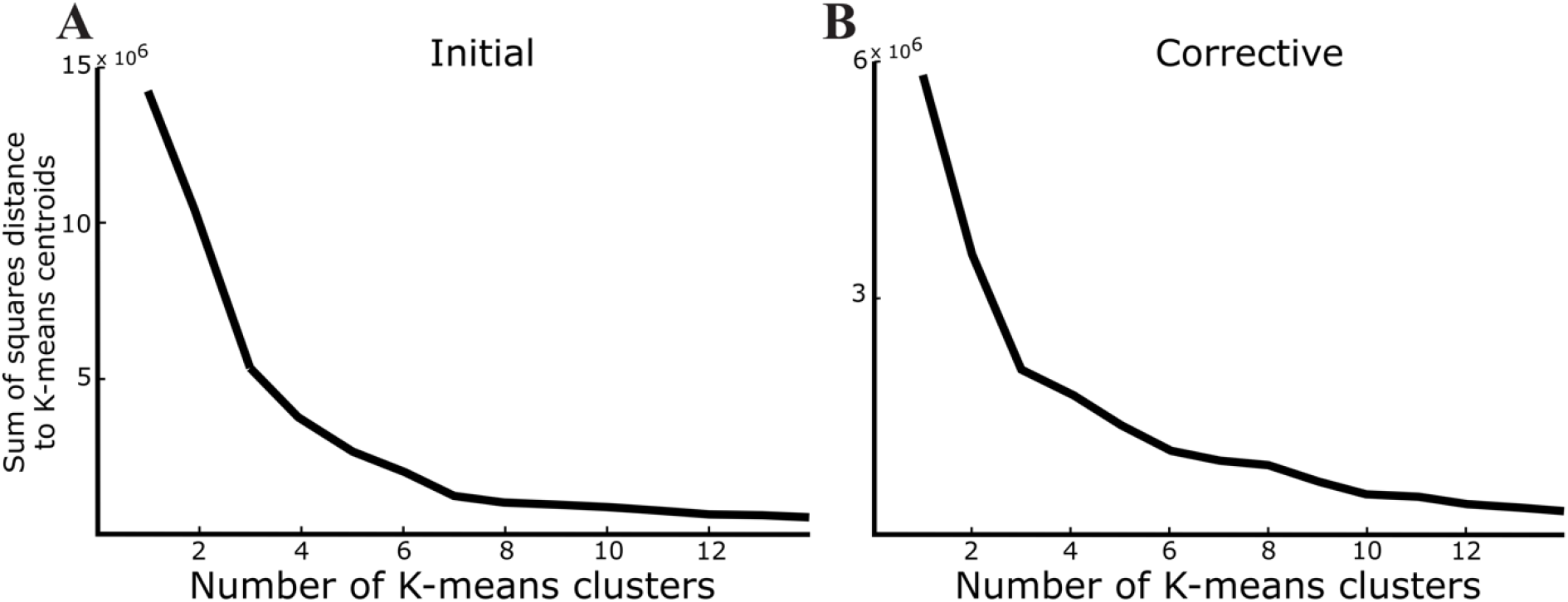
Elbow method for *k*-means to identify cluster number for the t-SNE performed on initial and corrective submovement LFADS factors. The *k*-means cluster number was chosen using the elbow method, visually inspecting the point where additional clusters didn’t significantly improve movements’ distance from the centroid. We identified 8 clusters for initial submovements (A) and 9 for corrective submovements (B).

A similar analysis was then performed on the corrective submovements. Fig. 5A shows a t-SNE plot of the corrective submovement LFADS factors. Although the corrective submovements had less distinct groups than initial in the t-SNE space, we again used *k*-means clustering to subdivide the neural data for display. We chose to label 9 groups for visualization of the kinematics and neural trajectories of submovements with similar neural features (Ext. Fig. 4-1). Fig. 5B shows the neural trajectory plot of corrective data in the same two principal components of the global LFADS factors space. For the corrective submovements, the average neural trajectories show smaller amplitude relative to initial trajectories. Additionally, the starting points of each cluster in corrective trajectories were relative to the ending point of initial trials in the LFADS factors space. Therefore, the smaller, corrective submovements do not begin at the same origin as the initial submovements and thus represent a different neural pattern than would be expected if a corrective submovement was encoded as just a smaller initial movement in a given direction. Similarly, we plotted the 9 *k*-means groups of submovements separately in the kinematic space to show the corresponding movement trajectories of this neural data (Fig. 5C). Seven of nine groups showed similar kinematic patterns covering many different starting positions in the upper portion of the workspace and were primarily grouped by reach direction. Two out of nine groups showed patterns covering only those movements occurring in the bottom positions of the workspace. In particular, the 6th and 7th group marked in the left yellow box and 8th and 9th group marked in the right blue box demonstrate good examples of 2 separate groups, each pair of groups with their own portion of the workspace and each group within the pair associated with different reach directions. A separate neural trajectory plot for the 4 highlighted groups was presented in Fig. 5D. Fig. 5E shows how the position in neural space could define the workspace when looking in the movement trajectory. Overall, the results suggest distinct domains or types of corrective submovements rather than a single direction- or position-based organization, demonstrating the encoding complexity of corrective submovements.

**Figure 5.**
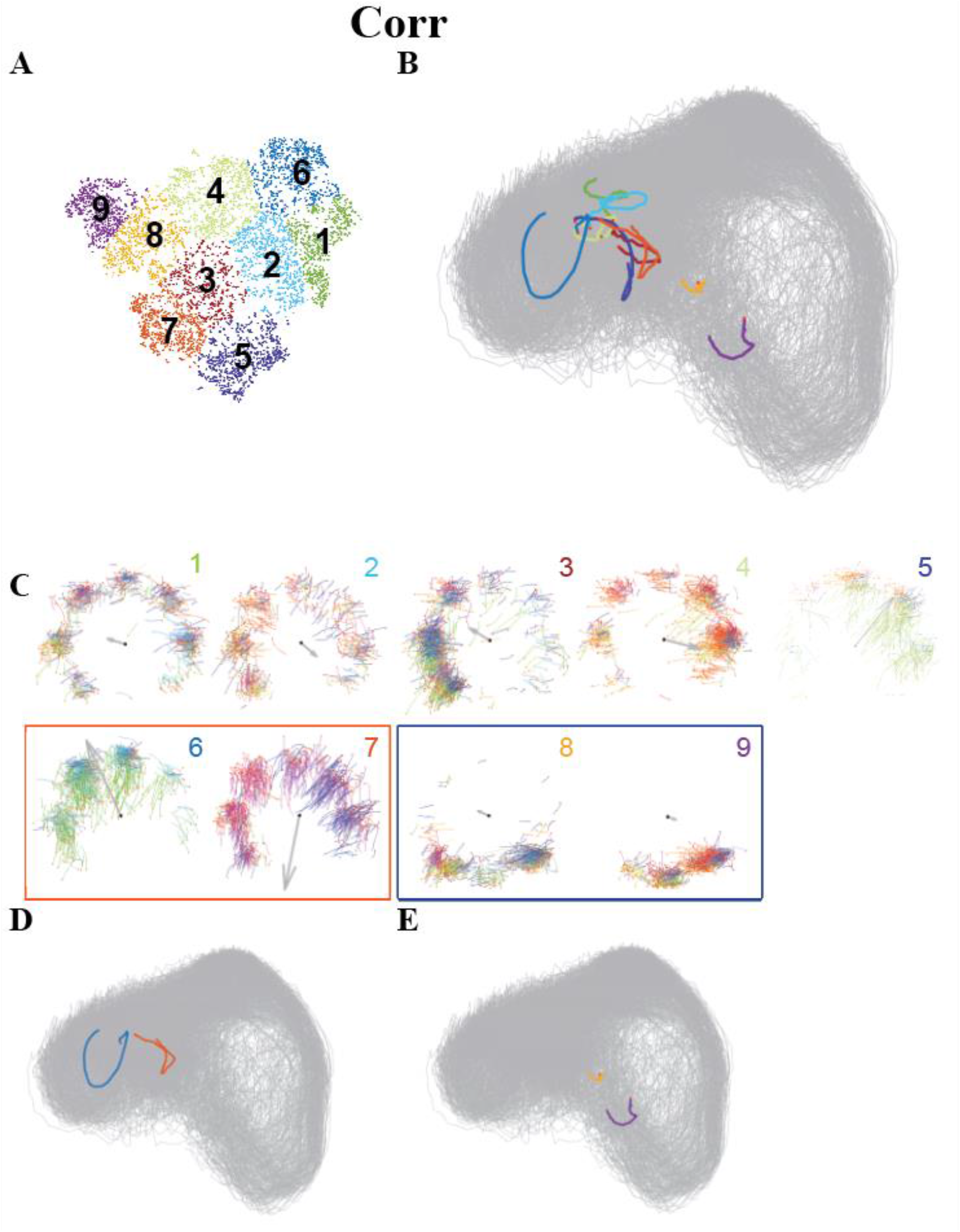
The identified organization of submovements in the neural space and their corresponding kinematics for corrective submovements. Similar plots as Fig 4 but for corrective submovements. A) t- SNE plots of initial and corrective submovements AutoLFADS factors, color coded by subgroup identified with *k*-means clustering. We selected the number of *k*-means groups to be 9 for corrective submovements using the elbow method shown in Figure 4-1. B) Neural trajectories of corrective submovements factors. Average neural trajectories are color coded based on the t-SNE subgroups. All neural trajectories are colored gray in the trajectory plot. C) Movement trajectories for each subgroup identified based on the neural activity. G,H) Neural trajectories plot for the 4 groups highlighted in dark yellow and blue rectangular. The two trajectories’ plots indicate different starting positions in neural space correlates with different movement trajectories in the workspace. Distinct neural dynamics between initial (Fig 4) and corrective (Fig 5) submovements were observed with corrective submovements occurring in a different location of the neural space. Also, the corrective submovements form subgroupings of movement that depend both on the current hand location following the initial reach and the subsequent movement direction.

To quantify this variation of starting neural states for corrective movements, the neural activity presented in Fig. 5B was examined for its dependence on both submovement type and cursor position at submovement onset. As shown in Fig. 6A & C, the neural activity for the initiation of corrective submovements is shifted from where it initiated for initial submovements. Additionally, the initial activity also depended on hand position in the work space as shown in Fig. 6B & D. The mean neural distance between initial and corrective submovements was 113% (monkey P) and 99% (Q) of the neural distance between the mean difference in neural activity for starting positions at opposite sides of the workspace. This indicates that the differences in neural activity at the onset of movement is explained by both the type of submovement—initial vs. corrective—and the hand position with approximately similar magnitude changes in firing rates for both.

**Fig 6.**
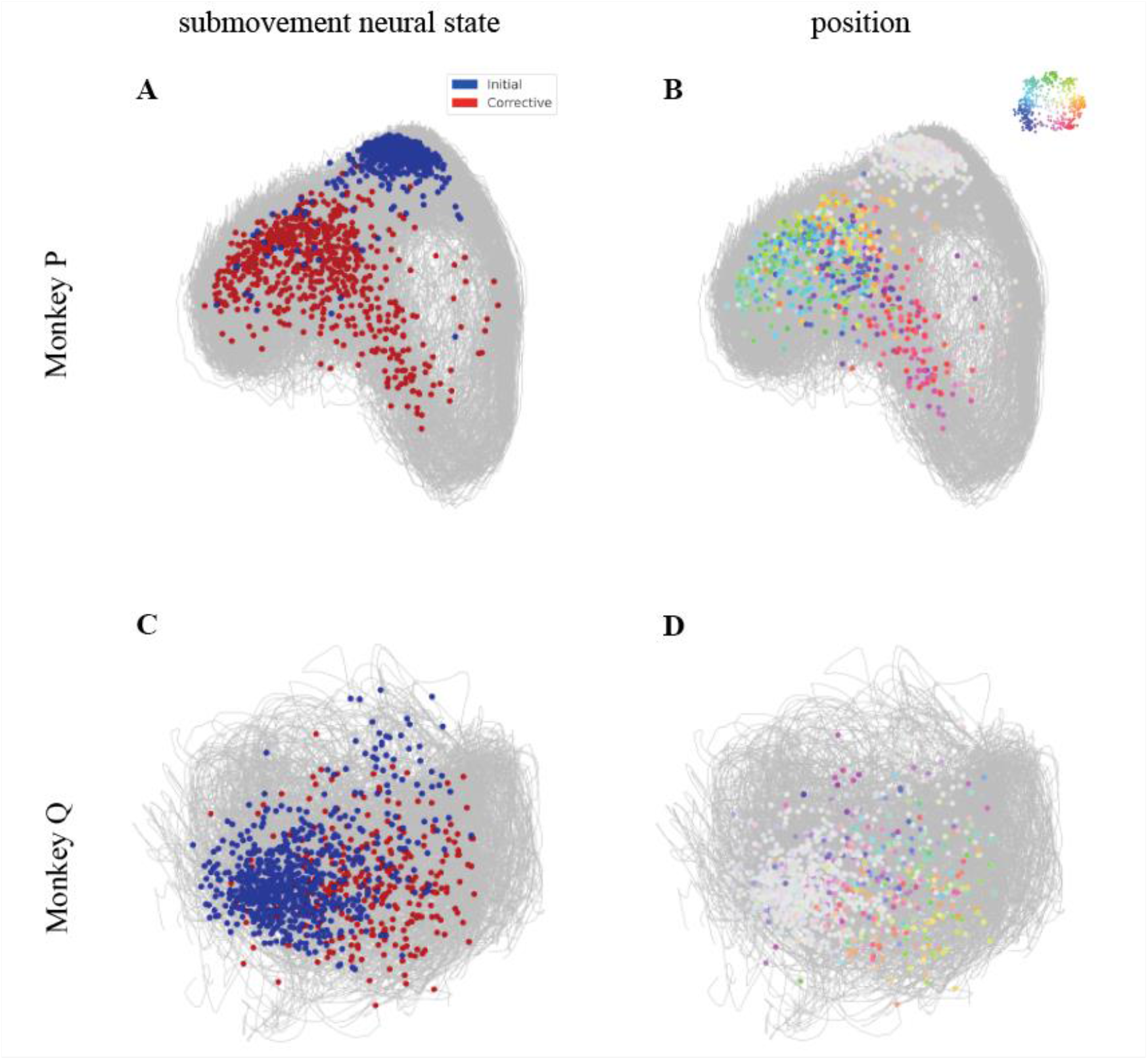
Neural states at initial time point labeled with submovement types and cursor positions. A) Initial (blue) and corrective (red) submovement trials color-coded in the latent neural space of monkey P at the initial time point. B) A total of 9 positions color-coded as shown in the color key in the latent neural space of monkey P at the initial time point. C) Initial (blue) and corrective (red) submovement trials color-coded in the latent neural space of monkey Q at the initial time point. D) A total of 9 positions color-coded as shown in the color key in the latent neural space of monkey Q at the initial time point. The average distance between initial and corrective submovement at initial time point (4.47 a.u. for monkey P, 3.48 a.u. for monkey Q) in comparison to the average distance between the maximum distance pairs (4 diagonal pairs across the workspace) in the workspace at initial time point (3.96 a.u. for monkey P, 3.51 a.u. for monkey Q) indicates the variability in initial neural state is not simply a position effect but rather a submovement effect.

### Decoding performance

We trained ridge regression decoders on the initial and corrective neural data to assess how well the brain activity could be decoded to predict movement and how well the decoders generalized between the two types of submovements. We trained decoders on LFADS factors of the two different types of submovements and applied the trained model to the other submovements. Fig. 7A-B shows decoders trained and tested either within or across submovement types. It also shows a generalized decoder trained on both submovement types and applied to either of the submovement types. Blue and red bars show the R² of the initial submovement velocity predictions and corrective submovement velocity predictions from the neural training data. For the decoder trained on initial submovement data, the predictions were poor for corrective submovements. Interestingly, the decoder for corrective submovements provided a reasonable prediction of initial movements. When a decoder was trained to fit all submovements combined, the decoding performance was worse for corrective submovements, indicating distinct patterns of neural activity with features unique to corrective movements. Thus, although AutoLFADS improved the R² of velocity predictions over smooth firing rates (Fig. 2), there remained issues with generalizability between the two movement state decoders.

**Figure 7.**
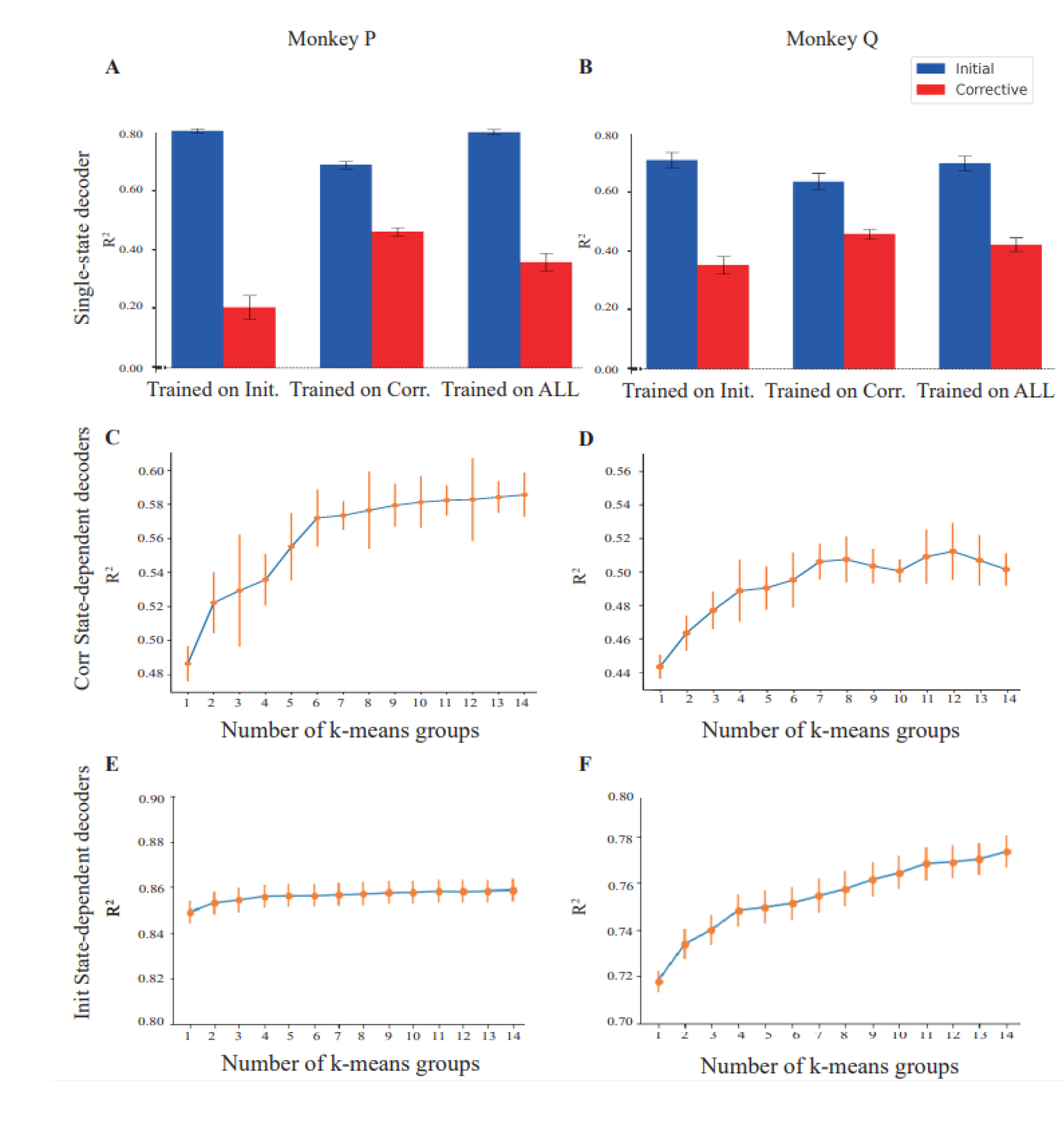
Linear decoders show improvement with state-dependent decoding in both monkeys. A-B) Decoding performance and generalizability for initial and corrective movements for monkey P (Fig. 6A) and monkey Q (Fig. 6B). Prediction accuracy of initial/corrective velocity from latent factors as measured with R^2^. The velocity of neural data is predicted for initial submovements in blue bars and corrective submovements in red bars using a ridge-regression linear neural decoder that was trained on initial (left), corrective (middle), or all (right) data. Showing poor generalization from initial to corrective submovements. Both monkeys showed significant decrease of decoding performances between trained on all and test on corrective vs. train and test on corrective, p-values < 0.05 with Mann-Whitney U test. C-F) Regression analysis using state-dependent decoders. Decoding performance on velocity and discrete neural state corrective movement decoders for monkey P (Fig. 6C) and monkey Q (Fig. 6D). E, F) Decoding performance on velocity and discrete state for initial submovements for monkey P (Fig. 6E) and monkey Q (Fig. 6F). Discrete state linear regression decoders were trained utilizing 1 (a single global decoder) to 14 separate states. For the discrete neural state decoder, each trial would be classified based on its neural state at the onset of the corrective submovement and then a linear prediction from a regression model specific to that subgroup was used to predict velocity. Error bars represent 95% confidence intervals for 1000 bootstrapped iterations. Showing subgrouping of corrective submovements improves predictions significantly compared to a single decoder over all corrective submovements. A comparison of predictions using clustering by hand position instead of the neural clustering shown here is provided in Figure 7-1. Neural clustering significantly improved compared to position for Monkey P and was similar for monkey Q.

Finally, we trained discrete-state linear decoders on the corrective movements as shown in Fig. 7C-D. This two-stage decoder i) divided the neural data into discrete states based on hand position and then ii) fit and evaluated a linear decoder only within that state. As the number of states increased, the decoding performance R² improved. For both animals, the performance plateaus around 7-10 states, inclusive of the 9 groups we labeled in Fig. 5A. We also trained the same discrete-state linear decoder on initial movements and observed less change in performance for increasing numbers of states (Fig. 7E-F). The improved performance with more decoder states indicates that the corrective submovement neural activity is more diverse and cannot be described well using a single linear decoder. Rather, identifying submovements with similar starting neural features provides a better prediction despite the resulting reduction in training dataset size within each state. This again demonstrates that distinct neural patterns may underlie individual subtypes of corrective movements that cannot be well-represented by a global encoding space defined by velocity or other kinematic features.

We also trained discrete-state linear decoders on the corrective movements by dividing the neural data into discrete states based on the starting hand position rather than the starting neural state. As described above, we tested how well velocity could be predicted for each number of states (Ext. Fig. 7-1). For Monkey P the decoder with neural clustering out-performed the decoder with position clustering, indicating discrete neural states contain more information than just hand position that is useful for decoding. For Monkey Q, the position effect appears slightly larger with no clear difference in decoding performance with neural or position clustering. This might be caused by the speed profile difference between the two monkeys as Monkey Q had more trials (13% of the total trials) with overlapping peak speeds between initial and corrective movements than did Monkey P (5% of the total trials) (Ext. Fig. 1-1). Overall, using the animal’s neural state at the start of each corrective movement appeared to include cursor position information for better reach predictions, with additional information beyond hand position being present for Monkey P.

**Figure 7-1.**
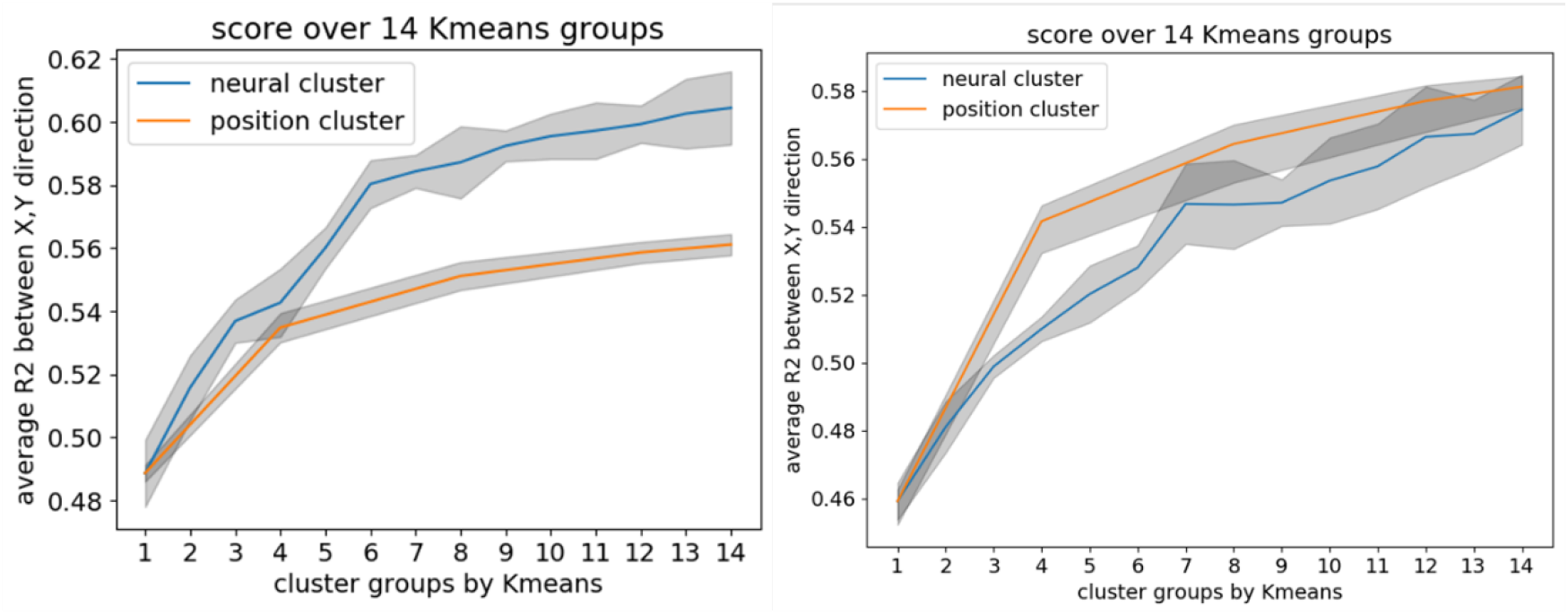
Comparison between state-dependent decoders trained to utilize either initial neural state or initial cursor position. The mean performance of the trained decoders for either neural states (blue) or initial positions (orange) over different numbers of clusters are shown. Shaded regions indicate the standard deviation of the performance across the decoders. For both the position and neural clusters, the decoder tests are generated by 5-fold cross-validation over each session.

## Discussion

Our results highlight that the neural encoding of planning and execution for corrective reaching movements is more diverse than projecting neural activity into a single linear plane or hyperplane for each movement feature such as velocity, force, or position. Previous studies have built linear models with time invariant instructed movements with a common origin in the workspace and produced reasonably accurate predictions (Moran and Schwartz, 1999; Serruya et al., 2002; Taylor et al., 2002; Santhanam et al., 2006; Wang et al., 2010; Gilja et al., 2012). These datasets are most similar to the initial movements in this study. However, the decoding performance of similar models was much worse for our self-generated corrective movements explored here. While some have previously reported the importance of neural fragments (Hatsopoulos et al., 2007), trajectories (Yu et al., 2007), or sequential reaches (Shanechi et al., 2012) for accurately representing neural encoding of reaching, the ability to organize this neural activity into the neural space on a single trial basis such as individually unique corrective movements has remained challenging.

### AutoLFADS provides a method to analyze consistent single-trial neural features across sessions

We have demonstrated that AutoLFADS, which had previously only been documented for applications to single datasets (Keshtkaran et al., 2022), can be used for “stitching” applications to extract generalized neural features across many recording sessions. AutoLFADS was useful for distinguishing unique neural features during the initial and corrective phases of precise reaching movements as corrective movements are rarely similar within individual sessions. Stitching with AutoLFADS enabled us to extract consistent neural dynamics that were more predictive of kinematics than smoothed spiking data (Fig. 2). Additionally, by utilizing t-SNE and subgrouping the neural latent factors, we could analyze these denoised neural features in a manner relatively agnostic to any underlying model of reach kinematics. That is, rather than searching for neural firing rates representing experimenter-defined kinematic features (e.g., with tools such as linear regression), we could examine trials when motor cortical activity was similar and then identify the commonalities and differences with regards to the corresponding kinematics of this similar-appearing neural activity. These results demonstrate that AutoLFADS is a powerful tool in evaluating high-dimensional neural populations across different sessions and during complex behaviors.

### Distinct neural features for initial and corrective phases of precise reaching

A previous study has shown uninstructed movements can dominate single-cell and population activity throughout the brain, outpacing task-related activity (Musall et al., 2019). Analysis of our own dataset showed a similar condition-independent cyclic neural trajectory pattern defining both initial and corrective submovements (Rouse et al., 2022). We now show here that when examining the neural space, while the neural trajectory contains the cyclic structure, the neural trajectories for corrective movements also lie in an entirely different portion of the neural space for some neural dimensions (Fig. 5B and Fig. 6). Unlike initial movements (Fig. 4A), latent factors of corrective movements cannot be clearly classified under a single 2D plane when multiple kinematic features define the movements being encoded (Fig. 4B). Notably, this difference in corrective movements could not be explained by differences in starting position in the workspace as would be expected in a linear velocity plus position model (Moran and Schwartz, 1999) as the neural differences between initial and corrective movements were as large as any neural differences due to hand position. Rather, the dominant neural feature was a small number of separate starting neural firing rates for most corrective submovements.

In addition to trajectory differences between submovement types, there were also differences in the ending positions of those trajectories, especially for initial movements. The divergence of initial movement neural trajectories (Fig. 4B) indicates potential posture information for different reaching targets in the high dimensional neural space. The neural trajectories for targets on the bottom-right and bottom appeared to propagate to a different location in the neural space. This may be explained by the constraints from the experimental setup where the monkeys are trained right-handed to acquire targets with the manipulandum, and more effort would be required to reach inward as this involves mostly flexion. Further experiments would be needed to quantify these posture effects.

### Corrective submovements cannot be generalized using a single decoder

When applying a single linear decoder, initial and corrective movements cannot both be decoded well. The corrective movements were especially poorly decoded by the initial movement decoder (Fig. 7A, 7C). We observed that the neural latent factors had large shifts for initial versus corrective movements, and corrective neural features were more diverse. Therefore, we built multiple state-dependent decoders on clusters of the neural data to examine whether neural features distinct for subtypes of movements helped describe the neural encoding of corrective movements.

Indeed, the corrective movements were not well organized into a single, linear encoding model in the neural space. When neural activity was decoded with only the most similar neural patterns in the neural space rather than fitting the neural features to a global space, decoding performance increased (Fig. 7B). The improvement continued to increase for up to at least 10 local subgroups. This suggests that the manifold for encoding corrective submovements is highly local with neural features not mapping neatly in a single, linear velocity space. Rather, idiosyncratic patterns of neural activity for subtypes of corrective movements appear to better define and predict the direction of a corrective movement. We hypothesize the motor cortex relies on a variety of corrective motor plans to generate a wide variety of corrective movements. The much higher-dimensional neural space of motor cortex would allow for a diverse set of patterns, all with convergent synaptic connections to the common lower-dimensional output of motor units.

### Future implications

Understanding the range of neural features that arise across submovements is important for optimizing brain-computer interfaces (BCIs). Historically, most BCI decoders for continuous control have assumed simple (e.g., linear, time-invariant) relationships between neural activity and intended movements (reviewed in (Pandarinath and Bensmaia, 2022)). More flexible decoding approaches, such as multi-state decoders (Williams et al., 2013; Sachs et al., 2016) and neural networks (Sussillo et al., 2012, 2016; Willsey et al., 2022; Deo et al., 2023) have introduced the ability to capture both nonlinear relationships between neural features and intended movements, and potentially any state-dependence in this relationship. Here we demonstrated that linear decoders that are localized to different regions of neural state space can improve decoding of single-trial movements. Helping to elucidate techniques that may allow flexible decoding strategies to improve BCI performance and can guide data collection and data augmentation strategies when training flexible decoders.

The control of a physical arm, as in our experiment, with more degrees-of-freedom than output dimensions, may differ from the control of a low-dimensional BCI cursor or limited degree-of-freedom robot; it may therefore face challenges in significantly improving predictions with fewer dimensions of control and less natural feedback. However, it also highlights that natural movement often relies on motor redundancy and a hypothesized uncontrolled manifold with variation along several degrees-of-freedom but precise movement along the variables critical to the task (Bernstein, 1967; Scholz and Schoner, 1999). Such concepts could influence future BCI design. These insights are also important when considering how to train and incorporate latent variable models, which have the potential to provide complementary improvements in BCI stability (Pandarinath et al., 2018a; Farshchian et al., 2019; Karpowicz et al., 2022; Ma et al., 2022).

With the help of AutoLFADS, we were able to better reveal the neural dynamics between initial and corrective submovements across sessions. Even with stitching, our dataset is limited as the motor errors were self-generated by the animal and resulted in impromptu corrective submovements on individual trials. The sample size for each corrective movement phase including correction start position and correction velocity vector is thus still unbalanced even after data aggregation. Also, the corrective movements were also almost all smaller magnitude than initial submovements and thus it is impossible to distinguish between the effect of correction itself versus making smaller instructed movements in the neural features of this dataset. More tasks that likely include not just user-generated errors but multiple instructions and online visual or mechanical perturbations are needed to further quantify all types of corrective movements. Another limitation is whether similar cognitive error signals persist when switching to imagined or attempted reaching tasks with no overt movement, as may be used in a BCI context. Future experiments may better define purely cognitive errors and overt motor errors and analyze whether their respective corrective neural activity share similar patterns. Future work is also likely needed to identify features that best separate the corrective submovements to build state-dependent decoders that can transition under different movement states to best predict the kinematics.

## Acknowledgements

We would like to thank Dr. Marc Schieber and his laboratory at the University of Rochester for their assistance with data collection.

## Funding

Emory Neuromodulation and Technology Innovation Center (ENTICe) NSF NCS 1835364

DARPA PA-18-02-04-INI-FP-021

NIH Eunice Kennedy Shriver NICHD K12HD073945 NIH-NINDS/OD DP2NS127291

NIH BRAIN/NIDA RF1 DA055667 the Alfred P. Sloan Foundation the Simons Foundation as part of the Simons-Emory International Consortium on Motor Control (CP), NIH NIBIB T32EB025816 (BMK).

HHS | NIH |NINDS R00NS101127 (AGR)

## Extended Data

